# Dopamine reward prediction error signal encodes effort cost as negative reward value

**DOI:** 10.64898/2026.11.11.765978

**Authors:** Mark Burrell, Alexandre Pastor-Bernier, Wolfram Schultz

## Abstract

All life must balance the economic problem of allocating scarce energy resources to maximize reward. Economic theory proposes that rational (reward-maximizing) decision makers balance the possible benefits from rewards against the cost of efforts for obtaining the rewards. Here we measured how the reward prediction error signal of monkey dopamine neurons reflects the negative value of effort cost (joystick resistance) and compared it with the dopamine signal for the positive value of reward. The animals repeatedly made binary choices between options varying in reward amount and effort to obtain the reward. Midbrain dopamine neurons signaled the negative value of effort cost both at effort-predicting stimuli (as reduction of reward-predicting excitation) and at experienced effort (as inhibition with more effort than predicted and as excitation with less effort). These effort signals contrasted with the opposite signal reflecting the positive value of reward relative to prediction (reward prediction error; RPE). Thus, dopamine neurons code effort cost as negative value opposite to the positive value of reward. In occurring with both stimuli and outcomes, this dopamine signal complies with the formalism of temporal difference reinforcement learning.

## Introduction

Material rewards provide molecules that are necessary for welfare and survival. Thus, rewards have value for biological organisms. However, rewards do usually not appear out of nowhere but require effort to acquire them. Biological effort requires expenditure of energy, and that energy requires metabolism of molecules such as carbohydrates, fats and proteins. Thus, effort removes critical molecules and thus has negative value for biological organisms that constitutes a cost and reduces the net benefit of rewards. Such cost can be tested in monkeys and rats as movement resistance or movement number required for obtaining a reward (Hosokawa et al. 2013; Kennerley & Wallis 2009; Day et al. 2010). Indeed, our preceding behavioral study in the same animals as presently used validated a subtractive effort model for reward choices in monkeys (Burrell et al. 2023), which applies also to choices in humans (Fehr & Goette 2007; Croxson et al. 2009) and monkeys in other laboratories (Hosokawa et al. 2013; Pasquereau & Turner 2013).

The question from these behavioral assessments arises how the negative value of effort cost is processed by reward neurons, in particular by the extensively studied dopamine neurons that signal reward prediction errors (RPEs). Dopamine neurons are excited by positive RPEs from better rewards than predicted and inhibited by negative RPEs from worse rewards than predicted. To test how dopamine neurons code effort cost, we defined effort prediction errors (EPEs) as the difference between experienced and predicted effort. If dopamine neurons were to code effort cost as negative reward value, we should see dopamine responses to effort that are opposite to reward: more effort than predicted should elicit negative EPEs and thus dopamine inhibitions, and less effort than predicted should elicit positive EPEs and dopamine excitations. As dopamine responses occur also to reward-predicting stimuli that constitute higher-order rewards according to the temporal difference reinforcement model, dopamine responses should reflect effort cost also with effort-predicting stimuli.

## Experimental Procedures

### Animals

The experiments were conducted on two purpose-bred male rhesus macaques (*Macaca mulatta*):

Monkey A weighing 14.5 kg, and Monkey B weighing 11.5 kg. The animals had been previously trained in a binary choice task in which we measured their effort cost functions (Burrell et al. 2023). The animal research, authorized by United Kingdom (UK) license PPL 70/8295, was ethically reviewed, approved, regulated and supervised by the following UK and University of Cambridge institutions and individuals: UK Home Office, implementing the Animals (Scientific Procedures) Act 1986, Amendment Regulations 2012, and represented by the local UK Home Office Inspector; the UK Animals in Science Committee; UK National Centre for Replacement, Refinement and Reduction of Animal Experiments (NC3Rs); the University of Cambridge Animal Welfare and Ethical Review Body (AWERB), the University’s Biomedical Service (UBS) Certificate Holder, Welfare Officer, Governance and Strategy Committee, Named Veterinary Surgeon (NVS) and Named Animal Care and Welfare Officer (NACWO).

The two monkeys were seated in an adjustable primate chair in the laboratory during data acquisition and released every day into their home cage that provided housing with one or several other monkeys. In the laboratory, the animals were trained to work for liquid rewards under mild fluid deprivation for a few hours during five or six days of each week. The totality of fluid was provided by performance in the behavioral task and by additional subsequent daily access to ensure the approved daily fluid requirements were met.

### Behavioral task and analysis

Habituation to behavioral testing and initial task training in the laboratory involved presentation of two-dimensional visual stimuli, performance of joystick movements using specified efforts, and delivery of drops of blackcurrant juice rewards over the course of > 5,000 trials over several days. Following this initial period of several months, each animal was trained and then tested in binary choices between two simultaneously presented options (Fig. 1A-C), as described in more detail for our previous behavioral experiment (Burrell et al. 2023).

**Figure 1.**
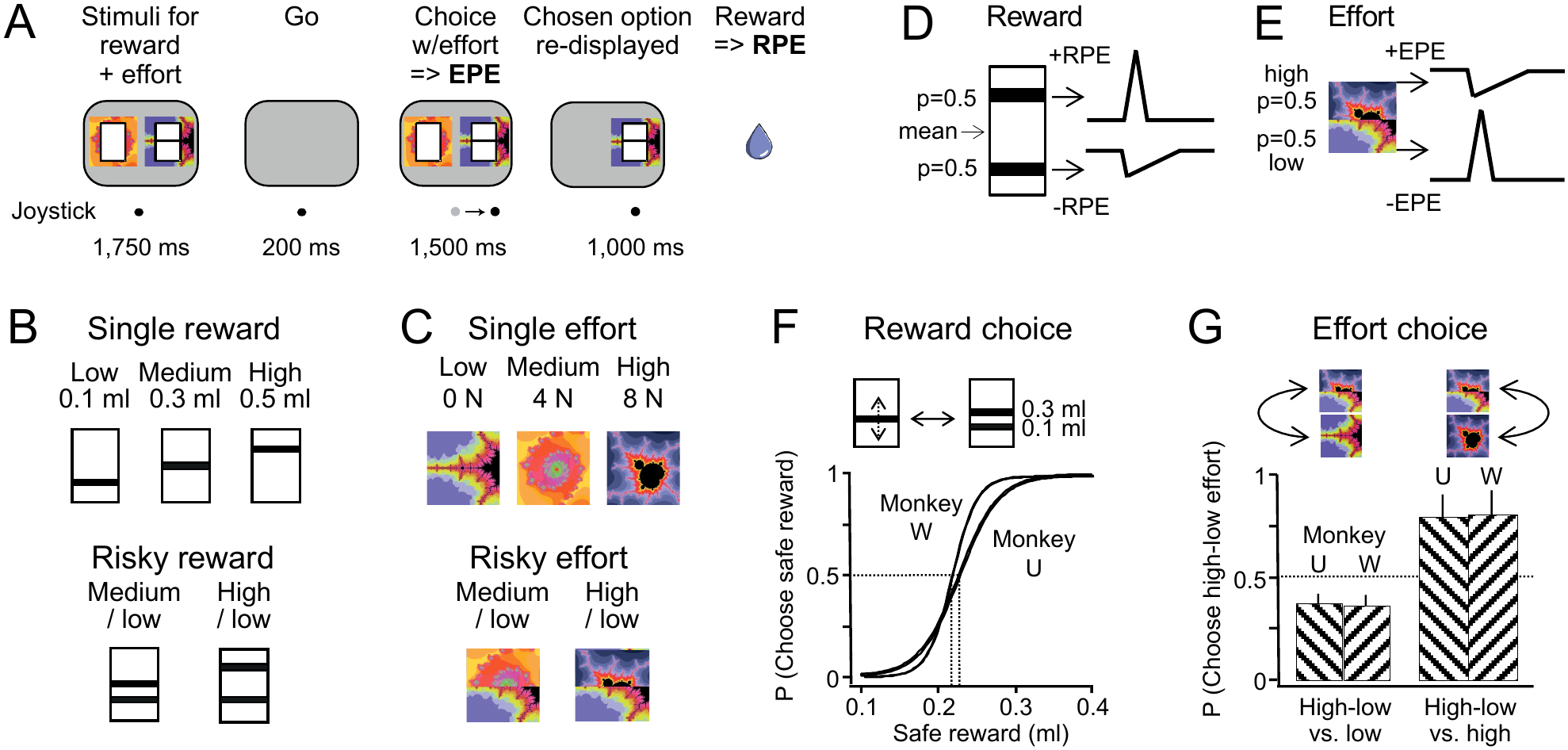
Behavioral task, stimuli, test design and behavior. (**A)** Event sequence within a trial (left to right): stimuli predicting specific reward amounts and joystick arm movement for left-right option choice. (B) Quantitative bars indicating single (‘safe’) juice reward amount (in ml; higher is more) and fractal stimuli indicating effort required for moving the joystick against a single friction level (in Newton). (C) Double-bar and spliced fractal stimuli indicating respective risky reward and effort (equiprobable and mutually exclusive reward amounts and effort levels (P = 0.5 each). (**D**) Design for eliciting reward prediction errors (RPE) by payout of one of two risky predicted reward amounts. (**E**) Design for eliciting effort prediction errors (EPE) by experience of one of two risky predicted efforts. (**F)** Psychophysical confirmation of meaningful reward choice (safe vs. risky options). (**G**) Preference hierarchy indicates meaningful effort choice: low effort is preferred to low-high effort mixture (P = 0.5 each effort), which is preferred to high effort. Note that all stimuli shown in panels B-F consisted of reward bars superimposed on effort fractals during the experiment, as shown in panel A.

Each trial started when the animal had moved a custom-made joystick to the center of the computer monitor in front of it (Fig. 1A). Then two composite images appeared that each consisted of one or two reward bars predicting specific amounts of juice reward. The reward bars allowed setting of reward amounts on a continuous scale (higher was more) for psychophysical testing (Fig. 1B, F). The reward bars were superimposed on fractal stimuli indicating the effort force required for moving the joystick against specific friction to obtain reward (for composite images, see Figs. 2A and 3A; for rationale and details, see Results section). The fractal stimuli indicated on a discrete scale the force required for obtaining the chosen option; the effort force was defined by the friction that needed to be overcome for moving the joystick for obtaining the reward. Subsequently, the stimuli disappeared, which reverted the monitor to grey and served as go signal for the choice, upon which the animal chose with the joystick one of the two stimuli and received the corresponding reward after a delay of 1 s. After the animal had made its choice, the stimulus of the chosen option reappeared to emphasize its association with the reward and effort. A variable intertrial intervall of 1–3 s with a blank screen ensued after each trial. Errors consisted of failure to center the joystick before stimulus presentation, maintain joystick contact, or make a choice within 2 seconds after the go stimulus. Errors resulted in trial repetition.

**Figure 2.**
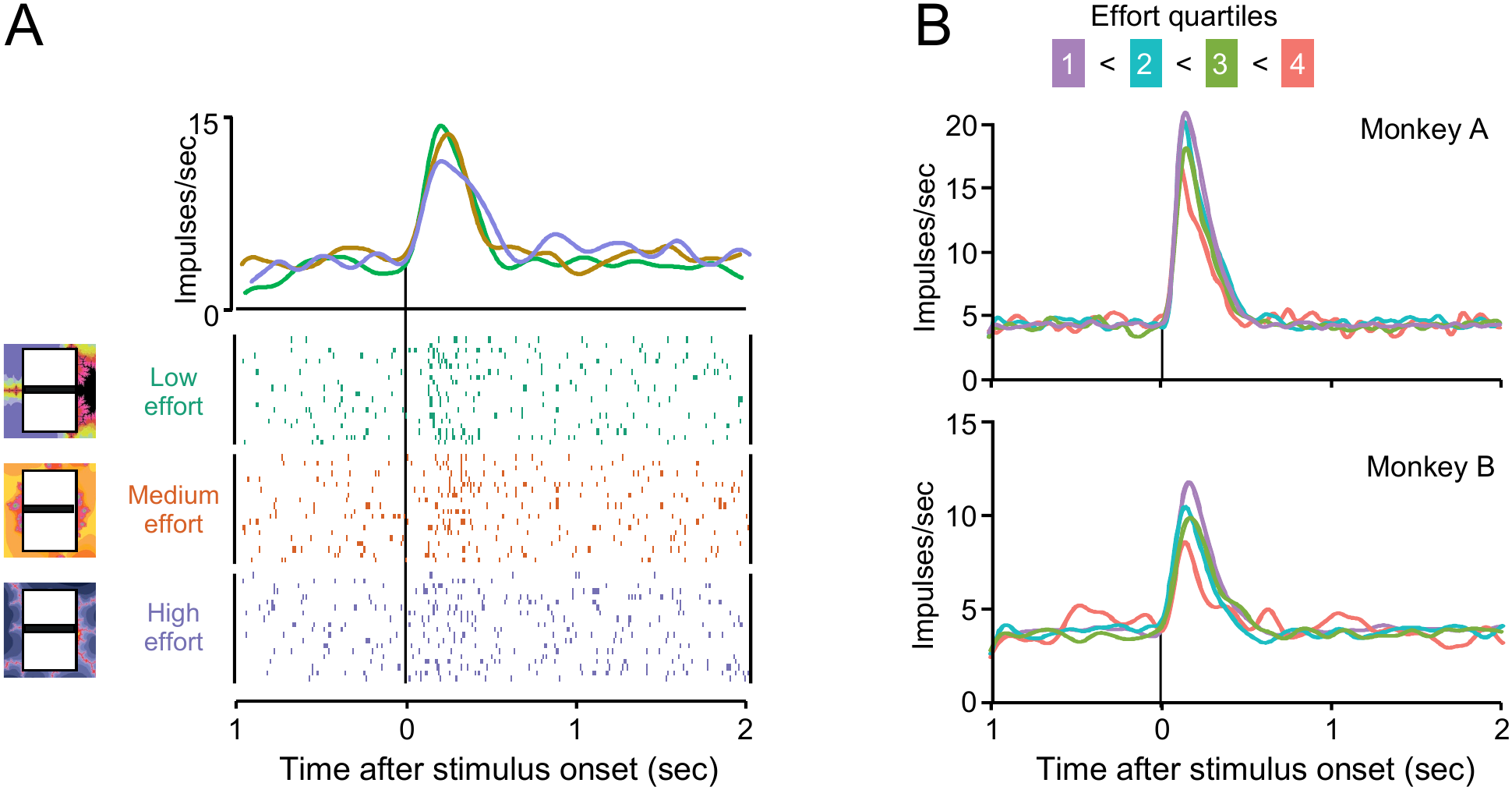
Decreasing dopamine responses to stimuli predicting increasing effort (with constant medium reward amount). (**A**) Single dopamine neuron. The combined reward-effort stimuli are shown on the left in corresponding colors. (**B**) Population responses in the two monkeys studied (four effort quartiles, from lowest to highest, 1 to 4). Top: 39 dopamine neurons, bottom: 68 dopamine neurons.

To determine the relevant variables the monkeys employed in their choice, we used a logistic model to estimate the probability of choosing a given side dependent on the option variables of reward and effort and on control variables (side, previous choice location):

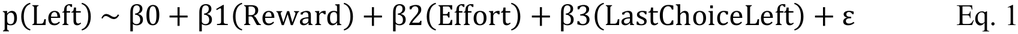

with p(Left) as probability of choosing left, β0 as side bias, β1 as sensitivity to reward (left minus right value), β2 as sensitivity to effort (left minus right), and β3 as tendency to choose the previously chosen side, with LastChoiceLeft as a binary variable that equals 1 if the left option was previously chosen, and 0 otherwise.

During the current experiment, and during its corresponding behavioral study (Burrell et al. 2023), Monkey A completed 159 weekday sessions with an average of 326 trials on each session (standard deviation (SD): 173 trials). Monkey B completed 122 sessions, averaging 298 trials each (SD: 138 trials).

### Surgical implantation

Each monkey was implanted with a titanium headpost (Crist Instruments Inc) for head fixation under general gas anesthesia and fully aseptic conditions. After the headpost had well integrated with the bone after several months, we implanted a recording chamber that permitted vertical access with single tungsten-in-glass microelectrodes (exposed tips of 2 x 10 micrometers) that were inserted into the brain for each recording session and withdrawn after each session. Chambers were centered on the skull using a stereotaxic head holder and a Kopf stereotaxic manipulator. After recovery from surgery, we drilled craniotomies above recording sites. Chambers were monitored and cleaned daily. Once the experiment was completed, recording sites were marked with electrolytic lesions (15–20 μA, 20–60 s). Subsequently, we sacrificed the animals by an overdose of sodium pentobarbital (90 mg/kg, IV) and subsequently perfused with 4% formalin in 0.1 M phosphate-buffered saline. We confirmed recording positions histologically from 40 μm coronal sections stained with cresyl violet.

### Ventral midbrain localization

Coordinates for recording sites in the ventral midbrain (substantia nigra pars compacta, ventral tegmental area) were determined using sagittal and coronal radiographs of the head in a stereotaxic frame and electrophysiological signatures. Specifically, the fully ketamine-anesthetized monkeys were placed in a stereotaxic frame and a cannula was inserted in the defined position of the horizontal (x-y) plane of the recording chamber. Bone features (interaural origin and clinoid process of sphenoid bone) served to determine the approximate anteroposterior and dorsoventral positions of familiar nuclei. Using these positions as anchors, stereotypical electrophysiological responses from the red nucleus and ventral posteromedial nucleus of the somatosensory thalamus (face area) guided the localization of recording sites of dopamine neurons in the ventral midbrain.

### Electrophysiology

We recorded the electrophysiological impulse activity of individual dopamine neurons at extracellular positions in the ventral midbrain during task performance using single moveable tungsten microelectrodes. Discharges from individual dopamine neurons were visualized on an oscilloscope and characterized by standard electrophysiological criteria: wide waveforms (> 1.8 ms), low baseline impulse rate (< 10 impulses/sec), and excitation by temporally unpredicted drops of juice reward relative to a pre-reward control period (positive RPE responses; p < 0.05, two-tailed Wilcoxon rank-sum test). Discharges of individual dopamine neurons were conventionally amplified, transformed into computer-compatible electrical signals via passage through an adjustable Schmitt-trigger under oscilloscope control, and displayed on-line during task performance as raster dot displays and peri-stimulus time histograms. Off-line statistical analyses were based on peri-stimulus time histograms and raster dot displays and employed nonparametric tests (Wilcoxon, Kruskal-Wallis) for comparison between pre-event and post-event time epochs or between behavioral situations.

### Results Experimental design

Each of two rhesus macaque monkeys (animals A and B) were seated for a few hours each weekday in a laboratory primate chair and used a joystick to choose one of two options that were simultaneously presented at the left and right in front of the animal (Fig. 1A). Each option was defined by a stimulus with one or two bars that predicted specific reward amounts. The bars were superimposed on a single or spliced fractal stimulus that predicted the effort required to obtain the reward. The choice served to monitor the animals’ understanding of the predicted reward and effort.

To test reward, reward amount was predicted by the vertical position of a bar inside a rectangle (higher was more; Fig. 1B top). Two bars predicted risky reward with two equiprobable and mutually exclusive reward amounts (P = 0.5 each reward; one, and only one, of which was paid out in any trial) (Fig. 1B bottom). The reward bars are shown in Fig. 1B in isolation for simplicity but were always overlaid onto the effort stimuli (see Fig. 1A). Rewards consisted of drops of blackcurrant juice with specific amounts (e.g. 0.1, 0.3, 0.5 ml).

To test effort, fractal stimuli predicted single effort levels that were required for overcoming the friction of a joystick for the left-right choice (Fig. 1C top). Spliced stimuli with two fractals predicted risky effort with two different, equiprobable and mutually exclusive efforts (P = 0.5 each effort; one, and only one, of which was applied in any trial) (Fig. 1C bottom). The effort stimuli are shown here in isolation but were always combined with the reward amount bars (see Fig. 1A). The three tested effort levels required forces of approximately 0 N (Newton), 4 N or 8 N force

To elicit RPEs, a risky stimulus with two reward bars served as reward predictor (Fig. 1D). After predictions stabilize with sufficient task experience, the risky stimulus predicts an intermediate reward value between the upper and lower amounts. Hence delivery of either reward amount elicits an RPE relative to the predicted intermediate value: the higher of the two amounts elicits a positive RPE and dopamine excitation, and the lower amount elicits a negative RPE and dopamine inhibition, as described before (Lak et al. 2014; Stauffer et al. 2014).

To elicit EPEs, a risky stimulus with a two-component spliced fractal image served as effort predictor (Fig. 1E). The animal found out during the movement which one of the two efforts was required for turning the joystick to get reward. After predictions stabilize with sufficient task experience, the risky fractal stimulus predicts an intermediate required effort between the upper and lower levels. Hence, in analogy to risky rewards, either of the efforts for the joystick turn differed from the predicted effort and elicited an EPE: experiencing the higher effort elicited a positive EPE, and experiencing the lower effort elicited a negative EPE. If, according to the hypothesis, effort has negative value, a negative EPE would indicate less negative value than predicted, equivalent to a positive RPE (‘less negative’ as double negation). By contrast, a positive EPE would indicate more negative value than predicted, equivalent to a negative RPE. If dopamine neurons process effort as negative reward value, they should be excited by negative EPEs and inhibited by positive EPEs.

### Behavior

In psychophysically controlled choices between a pseudorandomly varied single reward and a fixed risky reward (and equal effort), both animals showed monotonically increasing probability of choosing higher amounts of safe rewards against fixed risky alternatives (Fig. 1F; Eq. 1). In choices between a fixed effort (indicated by a single fractal) and a fixed risky effort (indicated by a spliced fractal image), both animals preferred risky low-high effort to safe high effort; the animals also preferred safe low effort to risky low-high effort (Fig. 1G). With these three fixed force levels, the animals’ choices complied with statewise dominance (lower effort is always equal to or better than higher effort), which is a minimal requirement for meaningful stimulus understanding and valuation. When reward and effort were tested together, both animals preferred any risky option to the least valued option (low reward, high effort). The observed effort choices allowed us to establish an ascending order of value of effort, from high safe effort (lowest value) via risky low-high effort (intermediate value) to low effort (highest value). These choices suggested that the tested monkeys valued effort inversely to reward.

To avoid confounds from the presentation of multiple stimuli and from choices during neuronal recordings, we set one of the two choice options to zero reward amount and highest effort. As a result, both animals almost exclusively chose the better option, i.e. non-zero amount and lowest effort (Monkey A: 99.2% of trials, Monkey B: 99.5% of trials). Thus, as dopamine neurons code chosen value (Morris et al. 2006; Lak et al. 2014; Ferrari-Toniolo et al. 2026), we assumed that these dopamine responses reflected the reward and effort of the better (chosen) option.

### Neuronal responses reflecting predicted reward value

A total of 170 dopamine neurons meeting the criteria described in Methods showed significant graded excitation to the stimuli predicting specific reward amounts (p < 0.05, two-tailed Kruskal-Wallis test; n = 88 Monkey A, n = 82 Monkey B), which confirms the reward sensitivity of the tested neurons and replicates their previously reported positive monotonic value coding (Tobler et al. 2005). Importantly, despite constant reward amount, these stimulus responses decreased with increasing effort, being lowest with highest effort, intermediate with intermediate effort, and highest with highest effort (Fig. 2A; green > red > blue).

The increases with decreasing effort were also observed in the dopamine population. Different from the initial behavioral tests, we varied the tested efforts occasionally during neuronal recordings for finer resolution. In these cases, we divided the effort into four quartiles (which often contained a mix of efforts varying within a small range) and averaged the responses of 107 dopamine neurons according to these four groups. As Fig. 2B shows, the dopamine population response to effort stimuli followed monotonically the inverse effort order in both monkeys, being lowest for the highest effort quartile, intermediate for the two intermediate effort quartiles in their order, and highest for the lowest quartile. Thus, the neuronal stimulus responses mirrored the negative effort value inferred from the behavioral preference for lower to higher efforts (Fig. 1G).

Thus, the prediction of the effort required to obtain a constant reward decreased dopamine responses to stimuli predicting this reward. The result suggests that dopamine neurons code effort as negative value. The issue of negative value coding of effort will now be tested separately from reward at the time of the experienced effort.

### Neuronal responses reflecting experienced effort

We first elicited bidirectional dopamine RPE signals to confirm the sensitivity of dopamine neurons to value prediction errors. Dual bar stimuli predicted two mutually exclusive equiprobable ‘risky’ reward amounts (probability P = 0.5 of each amount; for scheme, see Fig. 1D). Subsequent delivery of the larger of the two amounts elicited by definition a positive RPE and dopamine excitation, whereas the smaller amount elicited a negative RPE and dopamine inhibition, as documented before (Tobler et al. 2005; Lak et al. 2014; Stauffer et al. 2014). A total of 92% dopamine neurons showed such excitatory and inhibitory RPE responses at the time of reward delivery (Monkey A: 39 of 44 neurons; Monkey B: 61 of 65; p < 0.05; two-tailed Wilcoxon test), thus confirming the sensitivity of dopamine neurons to value prediction errors.

Next we tested dopamine neurons with bidirectional EPEs. To elicit EPEs, we presented the animal with two spliced fractal images after the go stimulus well before reward delivery (Fig. 1A); reward prediction and delivery were held constant. The two effort stimulus splices predicted two respective, mutually exclusive equiprobable efforts for overcoming the imposed joystick resistance (probability P = 0.5 of each effort; for scheme, see Fig. 1E). Following such a risky effort stimulus, the animal experienced the applied joystick resistance and thus the actual effort requirement at movement onset while making its choice. By definition, the larger of the two efforts would elicit a positive EPE, and the smaller effort would elicit a negative EPE (Fig. 1E). Thus, EPEs would occur before and distinct from reward delivery and thus distinct from RPEs, and any dopamine EPE response would occur separately from RPEs.

In almost all trials, the well-experienced animals moved the joystick continuously towards the chosen target (Monkey A: 98%, Monkey B: 97%). A typical dopamine neuron was inhibited by the positive EPE elicited by the higher of two risky efforts, and excited by the negative EPE elicited by the lower of two risky efforts. These responses were quite substantial with positive and negative EPEs elicited by the highest and the lowest experienced efforts (Fig. 3A, top) but more subtle or undetectable with intermediate EPEs elicited by intermediate efforts (middle) and absent with fully predicted efforts at any level (bottom).

**Figure 3.**
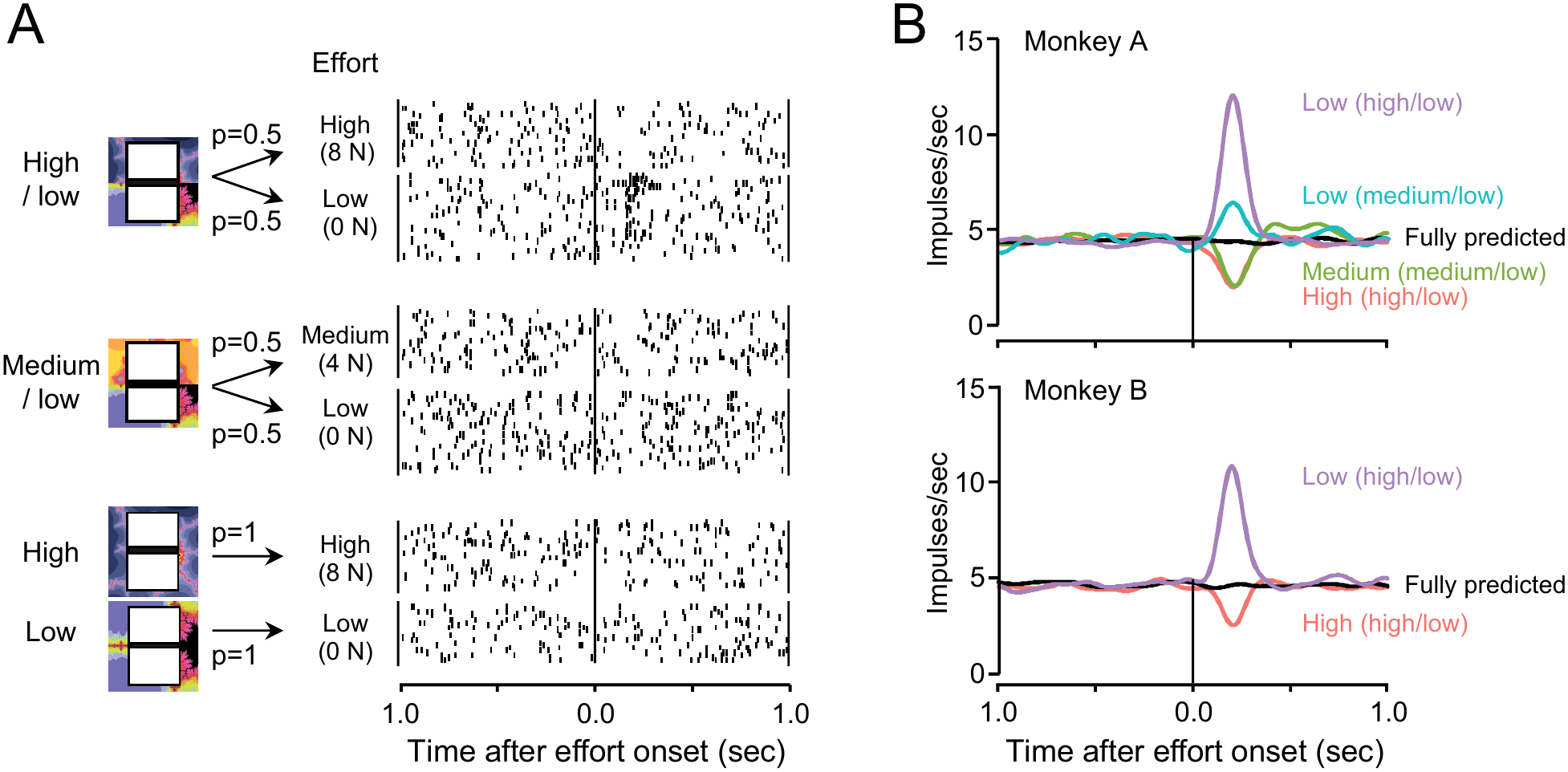
Dopamine responses to effort prediction errors (EPEs). (**A**) Single dopamine neuron. Top: after stimulus predicting high/low effort, the experienced high effort elicits a positive EPE and dopamine inhibition, whereas the low effort elicits a negative EPE and excitation. Middle: after medium/low prediction stimulus, the experienced medium effort elicits inhibition (no response to low effort, suggesting limited sensitivity). Bottom: no dopamine response to fully predicted effort. The combined reward and single/spliced effort stimuli are shown on the left. (**B**) EPE responses in dopamine population with analogous design as in A top and middle, demonstrating systematic processing of effort as negative value. Top: 44 dopamine neurons, bottom: 65 dopamine neurons. Predicted and delivered reward amount was constant at all effort levels.

Dopamine EPE responses became particularly prominent in the neuronal population. In both animals, experience of high effort (8 N) after equiprobable high-low effort prediction (8 N / 0 N, P = 0.5 each) elicited a positive EPE and substantial dopamine inhibition (Fig. 3B, orange). To the contrary, low effort (0 N) after the same high-low effort prediction elicited a negative EPE and substantial dopamine excitation (Fig. 3B, pink). Similarly, experience of medium effort (4 N) following the equiprobable medium-low effort prediction (4 N / 0 N, P = 0.5 each) elicited a positive EPE and a sizeable dopamine inhibition (Fig. 3B top, green), whereas low effort (0 N) elicited a negative EPE and a substantial dopamine excitation (cyan).

Importantly, the dopamine excitations just described elicited by the same experienced physical effort were larger with larger difference between experienced and predicted efforts that elicited larger negative EPEs (Fig. 3B top pink vs. cyan). By contrast, the stronger efforts failed to elicit significantly different dopamine EPE inhibitions between medium-low and high-low efforts (Fig. 3B top, green and orange), which may reflect a flooring effect of neuronal activity that cannot become negative, as noted before for RPEs (Stauffer et al. 2014).

Of 109 tested dopamine neurons, 79 neurons showed significantly different EPE responses between efforts fully predicted by a single fractal and risky efforts incompletely predicted by spliced fractals (Monkey A: 22 of 44 neurons; Monkey B: 57 of 65; p < 0.05; two-tailed Kruskal-Wallis test). Fully predicted efforts failed to elicit significant responses in individual dopamine neurons and in the whole tested dopamine population (Fig. 3B, black).

Thus, dopamine responses to EPEs varied inversely to dopamine responses to RPEs: positive EPEs from stronger-than-predicted efforts elicited dopamine inhibitions (similar to negative RPEs), and negative EPEs from weaker-than-predicted efforts elicited dopamine excitations (similar to positive RPEs). Importantly, these dopamine EPE responses occurred separately and independently from RPE responses and reward predictions, in contrast to the stimulus responses that reflected the combined value of reward and effort (Fig. 2). The bidirectional pure EPE responses are consistent with encoding effort cost as negative value opposite to reward: more effort is coded as less value and elicits dopamine inhibition, and less effort is coded as more value and elicits dopamine excitation.

## Discussion

This study suggests that dopamine neurons code effort as negative value and thus opposite to reward. The suggestion is based on two findings. First, dopamine responses to stimuli predicting fixed reward and variable effort decrease with increasing effort, indicating that effort reduces the combined value of stimuli predicting both reward and effort (Fig. 2). Second, dopamine responses to positive EPEs elicited by higher experienced effort than predicted consist of inhibitions, and dopamine responses to negative EPEs elicited by lower experienced effort than predicted consist of excitations (Fig. 3). Thus, the observed dopamine responses to EPEs are opposite to dopamine responses to RPEs, which reflects the general notion that effort has negative value opposite to the positive value of reward in these simple experiments. Thus, the notion of prediction error applies to dopamine signals for both positive and negative value for an organism.

The temporal difference reinforcement model account postulates that dopamine responses to stimuli reflect higher-order prediction errors relative to prediction from previous experience. The presently observed dopamine response to the combined reward- and effort-predicting stimuli likely reflect such prediction errors from additive combinations of RPEs and EPEs relative to previous predictions. As both the reward and the reward prediction were constant, the RPE was also constant. Thus, the decreasing dopamine stimulus response with increasing effort (Fig. 2) likely reflected an increasing dopamine inhibition from increasing EPE, similar to that seen at the time of the experienced effort (Fig. 3). The reduced dopamine stimulus response with higher effort would then be explained by a negative dopamine EPE signal, and the dopamine signals for effort cost can be incorporated into the prediction error framework of reinforcement learning.

The presently observed opposite prediction error coding between reward and effort may explain previously described dopamine responses during effortful behavior. In a fixed-ratio response schedule, stimuli predicting higher effort elicited lower dopamine concentration changes in the rat ventral striatum compared to stimuli predicting less effort (Day et al. 2010). Similarly, stimuli demanding higher effort for obtaining reward elicited lower excitations in midbrain dopamine neurons (Pasquereau & Turner 2013; Varazzani et al. 2015). Thus, these previously observed dopamine responses corresponded well to the present decrease of stimulus responses (Fig. 2). Besides being processed by dopamine neurons as negative value, the experienced effort also affects dopamine responses to reward itself: higher effort increases subsequent responses to reward-predicting stimuli and rewards in monkeys and mice (Tanaka et al. 2019, 2021; Eshel et al. 2024), which involves cholinergic action on dopamine axons (Touponse et al. 2026). In view of our present results, the likely explanation for these stronger dopamine excitations is the higher effort setting up a lower value prediction against which the subsequent constant reward-predicting stimuli and rewards elicit stronger positive RPEs. Thus, these earlier results are compatible with our notion of dopamine neurons coding the negative value of effort cost as inhibition.

Human neuroimaging studies suggest that the presently observed EPEs and RPEs originate in overlapping but spatially dissociable regions of the dopaminergic midbrain (Hauser et al. 2017).

The present data show that the same population of dopamine neurons codes both EPEs and RPEs, with opposite sign. This signed EPE coding contrasts with responses of noradrenergic neurons of the locus coeruleus, whose activation scales with the effort produced (Varazzani et al. 2015; Bornert & Bouret 2021) and whose outcome responses may signal attention-inducing salience and surprise irrespective of valence (Breton-Provencher et al. 2022). Such a noradrenergic mechanism may energize behavior to meet effort demands, whereas dopamine neurotransmission evaluates that demand relative to its prediction.

## Acknowledgements

We thank Aled David and Christina Thompson for animal and technical support, and Dr. Polly Taylor for implantation anesthesia. This study was supported by the Wellcome Trust (WT 095495, WT 204811, WT 206207), the European Research Council (ERC; 293549), and the National Institute of Mental Health (NIMH) Caltech Conte Center Grant P50MH094258. For the purpose of Open Access, the authors have applied a CC BY public copyright licence to any Author Accepted Manuscript version arising from this submission.

